# Myogenetic oligodeoxynucleotide restores differentiation and reverses inflammation of myoblasts aggravated by cancer-conditioned medium

**DOI:** 10.1101/2021.11.17.469038

**Authors:** Yuma Nihashi, Machi Yamamoto, Takeshi Shimosato, Tomohide Takaya

## Abstract

Cancer cachexia is characterized by irreversible muscle loss which is a critical factor in the prognosis of cancer patients. Myoblasts are myogenic precursor cells that are required to maintain skeletal muscle tissue. Previous studies have reported that cancer-released factors deteriorate myoblast differentiation, which is one of the causes of cachexia-associated muscle wasting. We recently identified the myogenetic oligodeoxynucleotide iSN04, which acts an anti-nucleolin aptamer and promotes myogenesis. The present study investigated the effects of iSN04 on human myoblasts exposed to conditioned medium (CM) of colon cancer cells. Cancer-CM impaired myogenic differentiation and myotube formation of myoblasts by upregulating the expression of inflammatory cytokines. iSN04 completely reversed cancer-CM-induced deteriorated myogenesis and inflammatory responses in myoblasts. Tumor necrosis factor-α (TNF-α), a representative cytokine present in cancer-CM, inhibited differentiation and induced inflammation of myoblasts, similar to cancer-CM. Pre-treatment with iSN04 reversed TNF-α-induced cachectic phenotypic features in myoblasts. These results indicate that iSN04 protects myoblasts against the effects of cancer-released factors and maintain their myogenic activity. This study provides a novel therapeutic strategy to prevent muscle loss associated with cancer cachexia.

## 1. Introduction

Cancer cachexia is a wasting syndrome characterized by an irreversible loss of adipose tissue and skeletal muscle mass along with anorexia, asthenia, and anemia [1]. Approximately 30-80% of cancer patients present with weight loss associated with chronic inflammation, and cachexia accounts for approximately 20% of cancer deaths [2]. As skeletal muscle is the primary organ for thermogenesis [3], energy storage [4], and insulin-responsive glucose uptake [5], a decrease in muscle mass disturbs systemic homeostasis and eventually increases the mortality risk of cancer patients. For instance, muscle wasting is a survival predictor for patients with metastatic colorectal cancer underlying chemotherapy [6]. Therefore, the prevention of muscle loss is a major problem in cancer therapy.

Myoblasts are the principal myogenic precursors involved in muscle formation. Myoblasts initially differentiate into myocytes, a process that is driven by myogenic transcription factors such as MyoD and myogenin [7]. Then, the myocytes fuse to form multinuclear myotubes by muscle-specific membrane proteins, myomaker and myomixer [8,9]. The factors released by the cancer cells have been reported to impair the myogenic program. Conditioned medium (CM) of colon cancer cells, including myostatin, promotes protein degradation in C2C12 murine myoblast cell line [10]. Prostate cancer-CM inhibits myogenic differentiation of murine myoblasts by upregulating CCAAT/enhancer binding protein β and interleukin (IL)-1β [11]. Hsp70 and Hsp90, associated with extracellular vesicles from lung and colon carcinoma cells, deteriorate murine myotube formation by disrupting catabolism [12]. Exosome miRNAs, miR-125b and miR-195a, in colon cancer-CM induce atrophy and apoptosis in murine myoblasts [13]. These cancerous factors have been confirmed to recapitulate muscle wasting in vivo [11–13]. Thus, exposure of myoblasts to the factors secreted by cancer cells is considered one of the causes of muscle loss, and myogenic ability of the myoblasts needs to be recovered to prevent cancer cachexia.

We recently reported that single-strand short DNAs named myogenetic oligodeoxynucleotides (myoDNs) promote the differentiation of myoblasts and rhabdomyosarcoma [14–17]. One of the myoDNs, iSN04, serves as an anti-nucleolin aptamer to increase p53 translation [14] and recovers myoblast differentiation attenuated by diabetes mellitus [15], demonstrating that iSN04 is a potential nucleic acid drug for disease-associated muscle wasting. The present study investigated whether iSN04 affects myogenesis and inflammation in human myoblasts cultured in colon cancer-CM mimicking cancer cachexia.

## 2. Materials and methods

### 2.1. Chemicals

Phosphorothioated iSN04 (5’-AGA TTA GGG TGA GGG TGA-3’) was synthesized and purified by HPLC (GeneDesign, Osaka, Japan) and dissolved in endotoxin-free water [14–17]. Recombinant mouse tumor necrosis factor-α (TNF-α) (Fujifilm Wako Chemicals, Osaka, Japan) was rehydrated in phosphate-buffered saline. Equal volumes of the solvent was used as the negative control.

### 2.2. CM preparation

The human colon fibroblast cell line CCD-18Co (CCD) (CRL-1459; ATCC, VA, USA) was used as a non-cancer cell line. The human colon adenocarcinoma cell line LoVo (IFO50067; JCRB Cell Bank, Osaka, Japan) and the human colon carcinoma cell line HCT-116 (HCT) (EC91091005-F0; ECACC, Salisbury, UK) were used as colon cancer cells. The cells were cultured in DMEM (Nacalai, Osaka, Japan) supplemented with 10% fetal bovine serum (GE Healthcare, UT, USA) and a mixture of 100 units/ml penicillin and 100 μg/ml of streptomycin (PS) (Nacalai) at 37°C under 5% CO_2_. The medium was replaced with non-supplemented DMEM when the cells became confluent. After 48 h, the medium was harvested as CM, filtered through 0.22-μm filters, and stored at −80°C.

### 2.3. Human myoblast culture

The commercially available human myoblast stock isolated from healthy subject (CC-2580; Lonza, MD, USA) was used [14–16,18]. The myoblasts were cultured on dishes coated with collagen type I-C (Cellmatrix; Nitta Gelatin, Osaka, Japan) at 37°C under 5% CO_2_. Undifferentiated myoblasts were maintained in Skeletal Muscle Growth Media-2 (CC-3245; Lonza). 7.5 × 10^4^ myoblasts were seeded on 30-mm dishes for immunocytochemistry, and 1.5 × 10^5^ myoblasts were seeded on 60-mm dishes for quantitative real-time PCR (qPCR). The next day, myogenic differentiation was induced by replacing the medium with differentiation medium (DM) consisting of DMEM supplemented with 2% horse serum (GE Healthcare) and PS. For CM experiments, CM was added at a final concentration of 30-50%. Non-supplemented DMEM was used as a negative control for CM. For TNF-α experiments, the myoblasts were pre-treated with 30 μM iSN04 for 3 h and then treated with 10 ng/ml TNF-α.

### 2.4. Immunocytochemistry

The myoblasts were fixed with 2% paraformaldehyde, permeabilized with 0.2% Triton X-100 (Nacalai), and immunostained with 0.5 μg/ml mouse monoclonal anti-myosin heavy chain (MHC) antibody (MF20; R&D Systems, MN, USA). 0.1 μg/ml of Alexa Fluor 488-conjugated donkey polyclonal anti-mouse IgG antibody (Jackson ImmunoResearch, PA, USA) was used as the secondary antibody. Cell nuclei were stained with DAPI (Nacalai). Fluorescent images were captured using EVOS FL Auto microscope (AMAFD1000; Thermo Fisher Scientific, MA, USA). The ratio of MHC^+^ cells was defined as the number of nuclei in all MHC^+^ cells divided by the total number of nuclei, and the fusion index was defined as the number of nuclei in multinuclear MHC^+^ myotubes divided by the total number of nuclei, which were calculated using the ImageJ software (National Institutes of Health, USA).

### 2.5. qPCR

Total RNA was isolated using NucleoSpin RNA Plus (Macherey-Nagel, Düren, Germany) and reverse transcribed using ReverTra Ace qPCR RT Master Mix (TOYOBO, Osaka, Japan). qPCR was performed using GoTaq qPCR Master Mix (Promega, WI, USA) with the StepOne Real-Time PCR System (Thermo Fisher Scientific). The amount of each transcript was normalized to that of the 3-monooxygenase/tryptophan 5-monooxygenase activation protein zeta gene (*YWHAZ*) [14]. The results are presented as fold-changes. Primer sequences (5’-3’) were as follows: myomaker (*MYMK*), CCC TGA TGC TAC GCT TCT TC and TCC AGC CTT CTT GTT GAC CT; myomixer (*MYMX*), ATC CAG CCA GAG ACT GAT TC and AGG ACA GCA GCA ATC GAA G. Primer sequences of IL-1β (*IL1B*), IL-6 (*IL6*), IL-8 (*CXCL8*), MyoD (*MYOD1*), myogenin (*MYOG*), myostatin (*MSTN*), nuclear factor-κB (NF-κB) p50 subunit (*NFKB1*), and TNF-α (*TNF*) were described previously [14,15].

### 2.6. Statistical analysis

Results are presented as the mean ± standard error. Statistical comparison between two groups was performed using unpaired two-tailed Student’s *t*-test and among multiple groups using Tukey-Kramer test following one-way analysis of variance. Statistical significance was set at *p* < 0.05.

## 3. Results

### 3.1. iSN04 recovers myoblast differentiation aggravated by cancer-CM

Cancer-CM has been reported to deteriorate myogenesis in murine myoblasts [11–13]. We first investigated the impact of colon cancer-CM on human myoblasts. Colon cancer cell lines LoVo and HCT were used to harvest cancer-CM. The colon fibroblast cell line CCD was used for control-CM. Non-supplemented DMEM served as the negative control. Human myoblasts were induced to differentiate using 50% CM for 48 h and immunostained for MHC (Fig. 1A). Both LoVo-CM and HCT-CM significantly decreased the ratio of MHC^+^ cells and multinuclear myotubes compared to that of non-supplemented DMEM and CCD-CM (Fig. 1B). This demonstrates that the colon cancer-released factors aggravated the myogenesis of human myoblasts. In the following experiments, LoVo, that shows undifferentiated signatures alike primary tumors [19], was used as a representative colon cancer cell line.

**Fig. 1.**
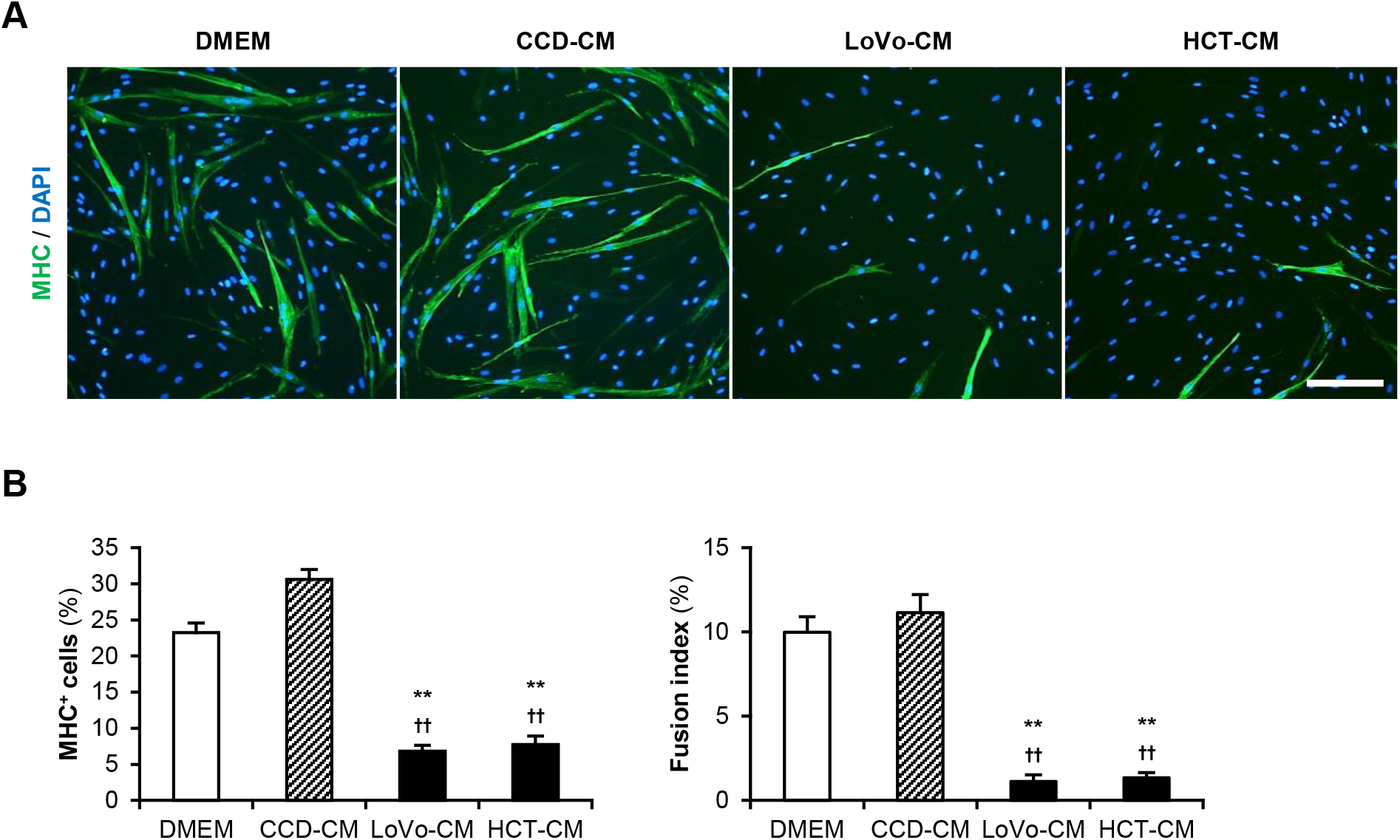
Cancer-CM deteriorate myoblast differentiation. (A) Representative immunofluorescent images of the myoblasts differentiated in DM with 50% CM for 48 h. Scale bar, 200 μm. (B) Quantification of the ratio of MHC^+^ cells and multinuclear myotubes. ** *p* < 0.01 vs DMEM; ^††^ *p* < 0.01 vs CCD-CM (Tukey-Kramer test). *n* = 6.

Next, we investigated whether iSN04 recovers the myogenic differentiation deteriorated by LoVo-CM. Human myoblasts were induced to differentiate using 30% CM and 30 μM iSN04 for 48 h (Fig. 2A). Consistent with the previous results, in the absence of iSN04, LoVo-CM significantly inhibited differentiation into MHC^+^ myocytes and myotubes compared to that of non-supplemented DMEM and CCD-CM. Interestingly, iSN04 markedly promoted myogenesis even in the presence of LoVo-CM (Fig. 2B). iSN04 completely restored the differentiation of the myoblasts exposed to LoVo-CM to the same extent as that to non-supplemented DMEM and CCD-CM. These results indicate that iSN04 enables myoblasts to differentiate even in the presence of cancer-released factors.

**Fig. 2.**
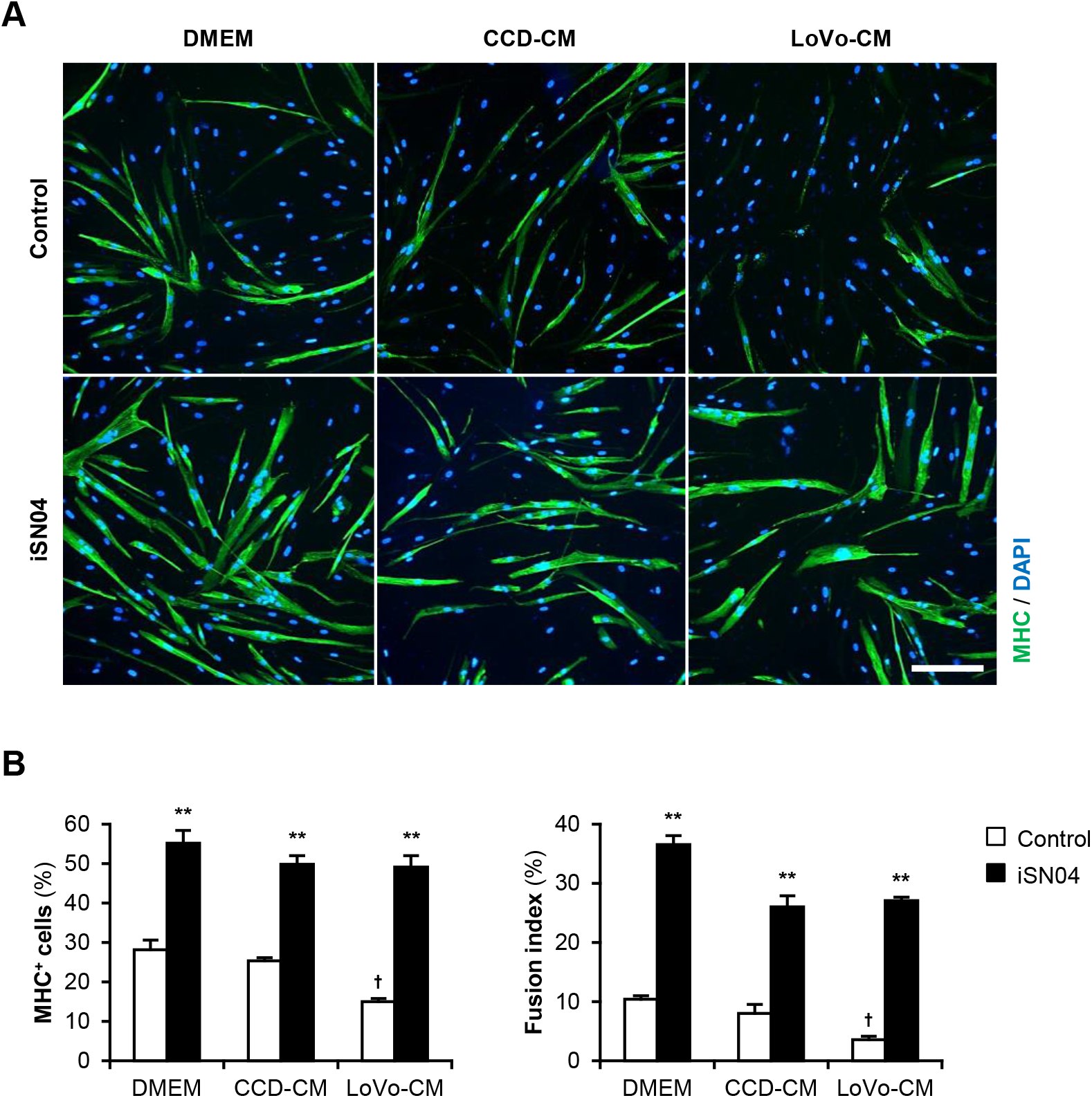
iSN04 recovers the myoblast differentiation deteriorated by cancer-CM. (A) Representative immunofluorescent images of the myoblasts differentiated in DM with 30% CM and 30 μM iSN04 for 48 h. Scale bar, 200 μm. (B) Quantification of the ratio of MHC^+^ cells and multinuclear myotubes. ** *p* < 0.01 vs control in each medium; ^†^ *p* < 0.05 vs DMEM-control (Tukey-Kramer test). *n* = 4.

### 3.2. iSN04 reverses gene expression in the myoblasts exposed to cancer-CM

To investigate the mechanism by which LoVo-CM deteriorates and iSN04 recovers myogenesis, the gene expression patterns in the myoblasts treated with LoVo-CM and iSN04 were quantified (Fig. 3). The mRNA levels of myogenic transcription factors, MyoD and myogenin, were significantly upregulated by iSN04 in LoVo-CM-treated myoblasts. Similarly, myomixer expression tended to increase under the same condition. iSN04-induced muscle gene expression is considered to cause the recovery of myogenesis in myoblasts cultured with LoVo-CM. However, it is still unclear how LoVo-CM impairs myoblast differentiation because LoVo-CM did not alter myogenic gene expression. qPCR revealed that LoVo-CM significantly elevated the mRNA levels of myostatin, IL-1β, and TNF-α in myoblasts, which are the cytokines inhibiting myogenesis [20–22]. iSN04 completely reversed LoVo-CM-induced transcription of the cytokines. These data demonstrate that iSN04 recovers myoblast differentiation by inducing myogenic genes and suppressing anti-myogenic cytokines.

**Fig. 3.**
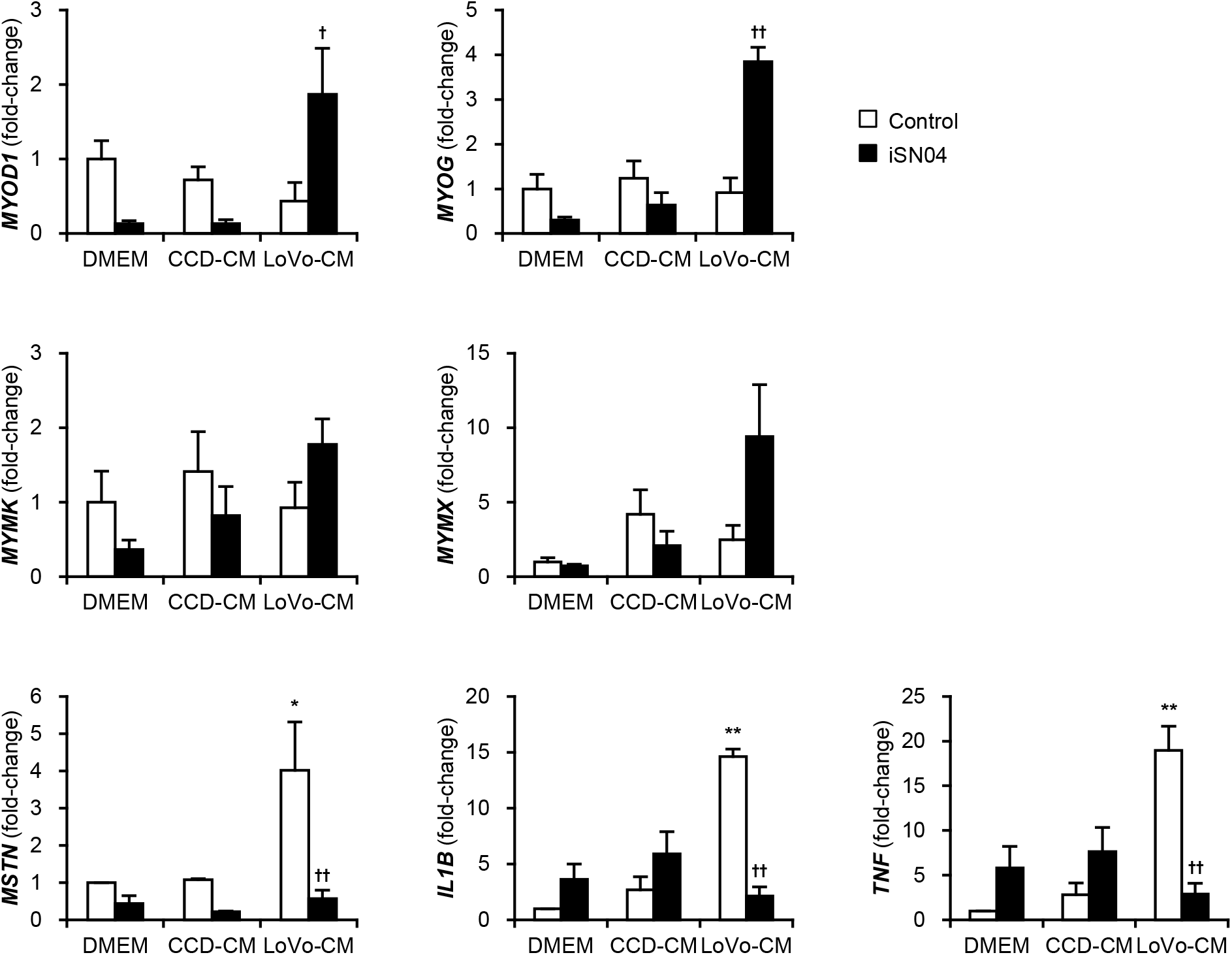
iSN04 reverses the gene expression patterns altered by cancer-CM. qPCR results of gene expression in the myoblasts differentiated in DM with 30% CM and 30 μM iSN04 for 24 h. Mean value of DMEM-control was set to 1.0 for each gene. * *p* < 0.05, ** *p* < 0.01 vs DMEM-control; ^†^ *p* < 0.05, ^††^ *p* < 0.01 vs LoVo-CM-control (Tukey-Kramer test). *n* = 3.

### 3.3. iSN04 reverses inflammation and attenuated differentiation of TNF-α-treated myoblasts

Cancer-CM contains a variety of cancer-released factors, which include multiple inflammatory cytokines [23]. qPCR indicated that LoVo expressed an inflammatory transcription factor, NF-κB, and its downstream cytokines, IL-1β, IL-8, and TNF-α, at significantly high levels compared to those in CCD (Fig. 4A). These cytokines present in LoVo-CM were speculated to initiate inflammatory responses and attenuate differentiation of myoblasts. TNF-α is a typical cytokine contained in cancer-CM [23] and has been reported to induce inflammation and impair myogenesis [20,24]. To investigate the effect of iSN04 on TNF-α-induced inflammation, human myoblasts were pre-treated with iSN04 for 3 h and then treated with 10 ng/ml TNF-α. The mRNA levels of IL-1β, IL-6, and IL-8 in the myoblasts were markedly increased by TNF-α, but significantly suppressed by pre-treatment with iSN04 (Fig. 4B). Correspondingly, pre-treatment with iSN04 completely reversed the attenuation of myoblast differentiation by TNF-α (Fig. 4C). These results demonstrate that iSN04 reverses inflammation and differentiation of myoblasts deteriorated by TNF-α.

**Fig. 4.**
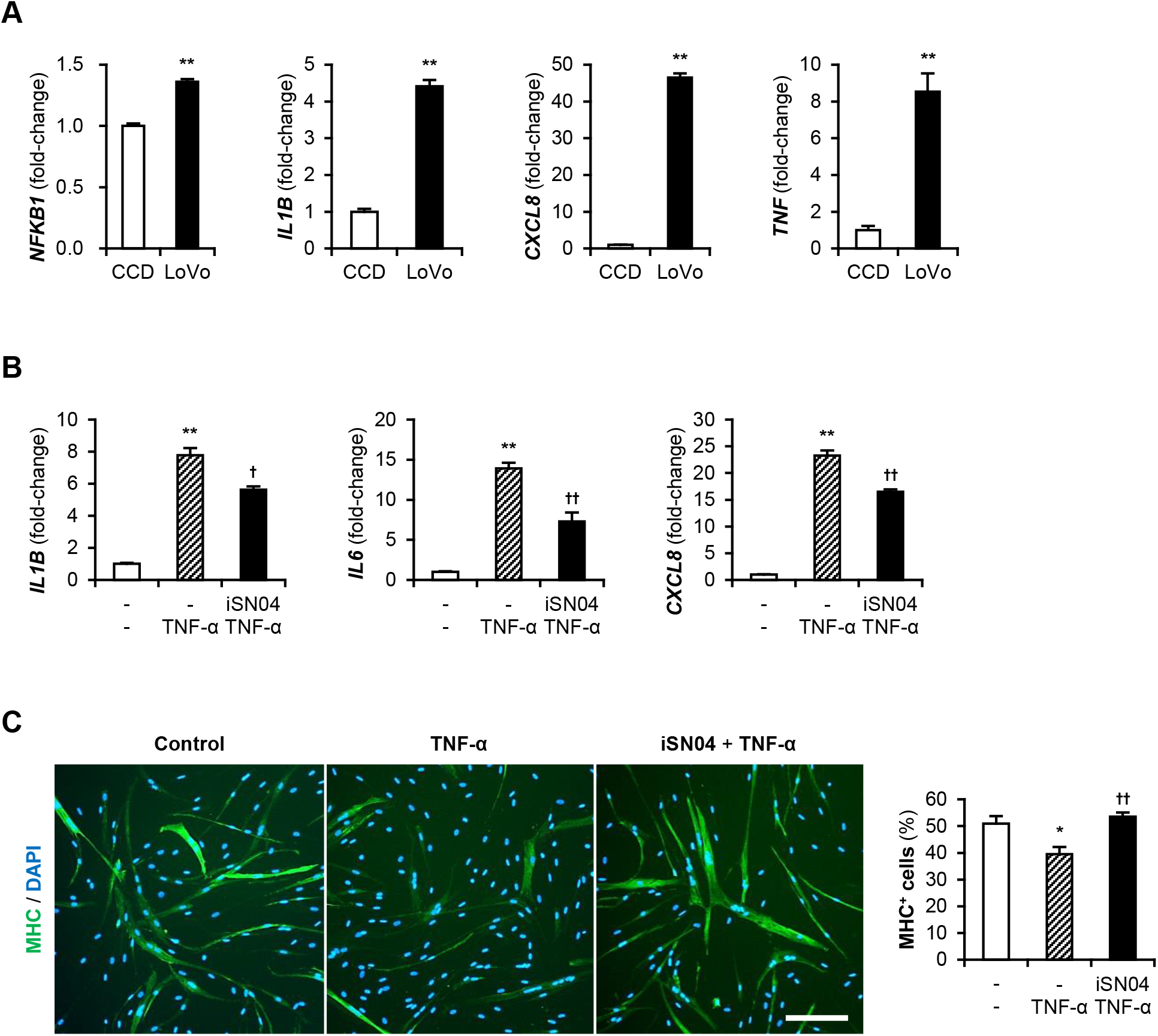
iSN04 reverses inflammation and restores differentiation of TNF-α-treated myoblasts. (A) qPCR results of gene expression in CCD and LoVo. ** *p* < 0.01 vs CCD (Student’s *t*-test). *n* = 3. (B) qPCR results of gene expression in myoblasts pre-treated with 30 μM iSN04 for 3 h and then treated with 10 ng/ml TNF-α for 1 h. ** *p* < 0.01 vs control; ^†^ *p* < 0.05, ^††^ *p* < 0.01 vs TNF-α (Tukey-Kramer test). *n* = 3. (C) Representative immunofluorescent images of myoblasts pre-treated with 30 μM iSN04 for 3 h and then treated with 10 ng/ml TNF-α in DM for 48 h. Scale bar, 200 μm. The ratio of MHC^+^ cells was quantified. * *p* < 0.05 vs control; ^††^ *p* < 0.01 vs TNF-α (Tukey-Kramer test). *n* = 5.

## 4. Discussion

The present study indicated that iSN04 rescued myogenic differentiation and reversed inflammatory responses in human myoblasts exposed to colon cancer-CM. Cancer-released factors, including cytokines and miRNAs, have been reported to impair myogenesis, one of the mechanisms of cachexia-associated muscle loss [11–13]. In this study, the colon cancer cell line LoVo showed high-level expression of IL-1β, IL-8, and TNF-α. As previously reported, TNF-α disturbed myoblast differentiation and evoked inflammatory responses, however, iSN04 successfully reversed TNF-α-induced cachectic phenotypic features in myoblasts. The protective effect of iSN04 against cancer secretions is expected to prevent deteriorated differentiation and inflammation in myoblasts of cancer patients. Myogenetic and anti-inflammatory effects of iSN04 have also been reported in diabetic human myoblasts [15]. These studies support that iSN04 can be a potential nucleic acid drug to prevent disease-associated muscle wasting along with systemic chronic inflammation such as cancer cachexia.

It is well known that inflammatory cytokines, particularly TNF-α, activate NF-κB, which not only enhances inflammatory gene expression but also impairs myogenic differentiation of the myoblasts [20,24]. Myostatin is a downstream of NF-κB [25] and a critical inhibitor of myogenesis [21]. As iSN04 reversed cancer-CM-induced upregulation of NF-κB-regulated genes (myostatin, IL-1β, and TNF-α), iSN04 is predicted to interfere with the NF-κB signaling pathway. iSN04 serves as an anti-nucleolin aptamer [14].

Nucleolin mediates NF-κB-dependent expression of IL-1β and TNF-α in monocytes [26]. AS1411, another anti-nucleolin aptamer that promotes myogenesis as well as iSN04 [14], forms a complex with nucleolin and NF-κB essential modulator (NEMO) to block the NF-κB signaling, and eventually suppresses TNF-α-induced inflammation in cancer cells [27]. Although the anti-inflammatory mechanism of iSN04 is speculated to be identical to that of AS1411, this needs to be further investigated using myoblasts.

In myoblasts, NF-κB downregulates the myogenic transcription factors, MyoD and myogenin [20,24,28]. The present study indicated that iSN04 significantly induces MyoD and myogenin in myoblasts cultured with cancer-CM. In our previous study, we demonstrated that iSN04 upregulates MyoD and myogenin expression by improving p53 protein level in myoblasts [14]. Nucleolin binds to the 5’ untranslated region of p53 mRNA and interferes with its translation [29,30], however, antagonizing nucleolin by iSN04 recovers p53 translation [14]. Numerous studies have reported that p53 promotes myogenesis [31,32], consequently, iSN04 accelerates myoblast differentiation by activating the p53 signaling pathway [14]. Intriguingly, NF-κB and p53 antagonize each other’s activity by competing with p300 [33]. This might explain why iSN04 upregulated myogenic gene expression even in the presence of cancer-CM.

The present study revealed the dual role of iSN04, myogenetic activity and anti-inflammatory effect, that restores the myoblasts exposed to cancer-CM. iSN04 is therefore anticipated to maintain myoblast activity in cancer patients by promoting myogenesis and suppressing inflammation, and eventually prevent cachectic muscle loss. The effects of iSN04 need to be studied in vivo using animal models to establish a novel therapeutic strategy for cancer cachexia.

## Author contributions

TT and YN designed the study and wrote the manuscript. YN and MY performed experiments and analyses. TS designed iSN04.

## Acknowledgments

This study was supported in part by a Grant-in-Aid from The Japan Society for the Promotion of Science (19K05948) to TT.

